# A methodology for unsupervised clustering using iterative pruning to capture fine-scale structure

**DOI:** 10.1101/234989

**Authors:** Kridsadakorn Chaichoompu, Fentaw Abegaz Yazew, Sissades Tongsima, Philip James Shaw, Anavaj Sakuntabhai, Bruno Cavadas, Luísa Pereira, Kristel Van Steen

## Abstract

SNP-based information is used in several existing clustering methods to detect shared genetic ancestry or to identify population substructure. Here, we present a methodology for unsupervised clustering using iterative pruning to capture fine-scale structure called IPCAPS. Our method supports ordinal data which can be applied directly to SNP data to identify fine-scale population structure. We compare our method to existing tools for detecting fine-scale structure via simulations. The simulated data do not take into account haplotype information, therefore all markers are independent. Although haplotypes may be more informative than SNPs, especially in fine-scale detection analyses, the haplotype inference process often remains too computationally intensive. Therefore, our strategy has been to restrict attention to SNPs and to investigate the scale of the structure we are able to detect with them. We show that the experimental results in simulated data can be highly accurate and an improvement to existing tools. We are convinced that our method has a potential to detect fine-scale structure.

## INTRODUCTION

Single Nucleotide Polymorphisms (SNPs) can be used to identify population substructure, but resolving complex substructures remains challenging [1]. Owing to the relatively low information load carried by single SNPs, usually thousands of them are needed to generate sufficient power for effective resolution of population strata due to shared genetic ancestry [2]. In practice with high-density genome-wide SNP datasets, linkage disequilibrium (LD) and haplotype patterns are likely to exist. These can be exploited for the inference of population structure [3], although exploiting haplotype patterns still comes with a high computational burden. Also, although removing LD by pruning strategies can eliminate some spurious substructure patterns [4], it may limit our ability to identify subtle subgroupings.

The identification of substructure in a study sample of healthy controls or patients is in essence a clustering problem. Several clustering algorithms exist, each leading to different clustering methods. Conventional population structure analyses use Bayesian statistics to show relationships amongst individuals in terms of their so-called admixture profiling, where individuals are clustered by using ratios of ancestral components (see also [5]). The iterative pruning Principal Component Analysis (ipPCA) approach differs from this paradigm as it assigns individuals to subpopulations without making assumptions of population ancestry [6].

At the heart of ipPCA lies performing PCA with genotype data, similar to EIGENSTRAT [4]. If substructure exists in a principal component (PC) space (ascertained using, for instance, Tracy-Widom statistics [6], or the EigenDev heuristic [7]), individuals are assigned into one of two clusters using a 2-means algorithm for which cluster centers are initialized with a fuzzy c-means algorithm. The test for substructure and clustering is performed iteratively on nested datasets until no further substructure is detected, i.e. until a stopping criterion is satisfied. The method is visualized in Figure 1. The software developed to perform ipPCA has some shortcomings though. Notably, it is limited to a MATLAB environment, which is not freely available. Also, outliers can severely disturb the clustering analysis. These limitations are addressed in IPCAPS, which improves the power of fine-scale population structure, while appropriately identifying and handling outliers.

**Figure 1.**
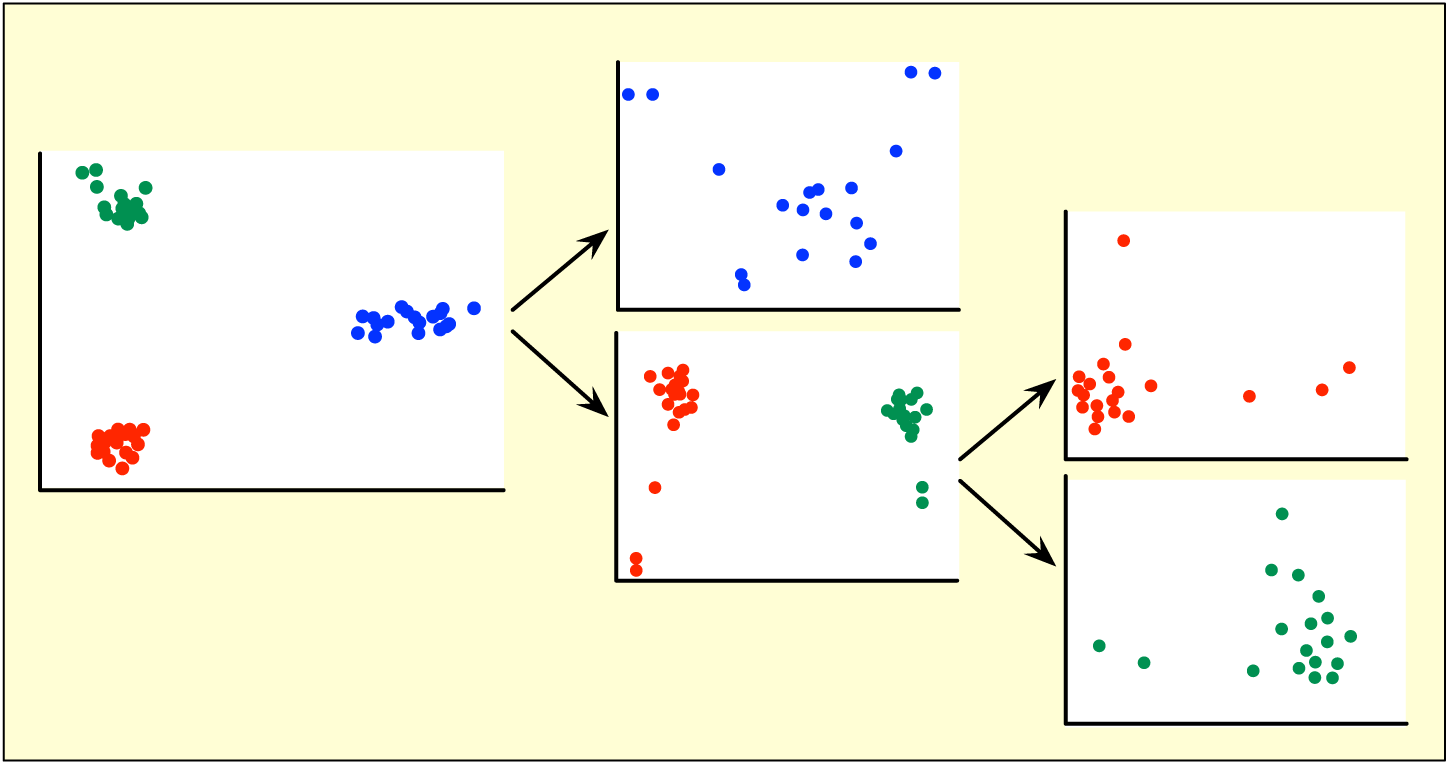
Illustration to show the iterative pruning process in ipPCA.

## IPCAPS METHODOLOGY

The IPCAPS methodology involves unsupervised clustering to identify sub-populations (the implementation of IPCAPS in R, see [8]). IPCAPs is a PCA-based method; therefore, the data quality control (QC) process for IPCAPS analysis is similar to the one typically adopted for classic PCA. In practice, data QC includes missing genotype filtering (missingness < 0.02), Hardy-Weinberg equilibrium (HWE) testing (p<0.001), and linkage disequilibrium (LD) pruning (r^2^<0.2 [9]). Once the input data are cleaned, a data matrix is constructed. The rows of this data matrix represent individuals, and the columns represent SNPs. SNPs are encoded as 0, 1, and 2, reflecting the number of minor alleles present at the corresponding loci. As a consequence, the encoded data matrix contains numeric values and has an order, which is suitable for PCA. The data matrix is subsequently normalized by a zero-mean and unit variance procedure. In case that all individuals contain only a single genotype at some loci, normalized value is zero representing no variation. This commonly happens for common alleles. Furthermore, each missing genotype is replaced by the most common value [10].

In particular, after a pre-analysis step involving data quality control, the major steps of the IPCAPS methodology are shown in Figure 2 and described below:

**Figure 2.**
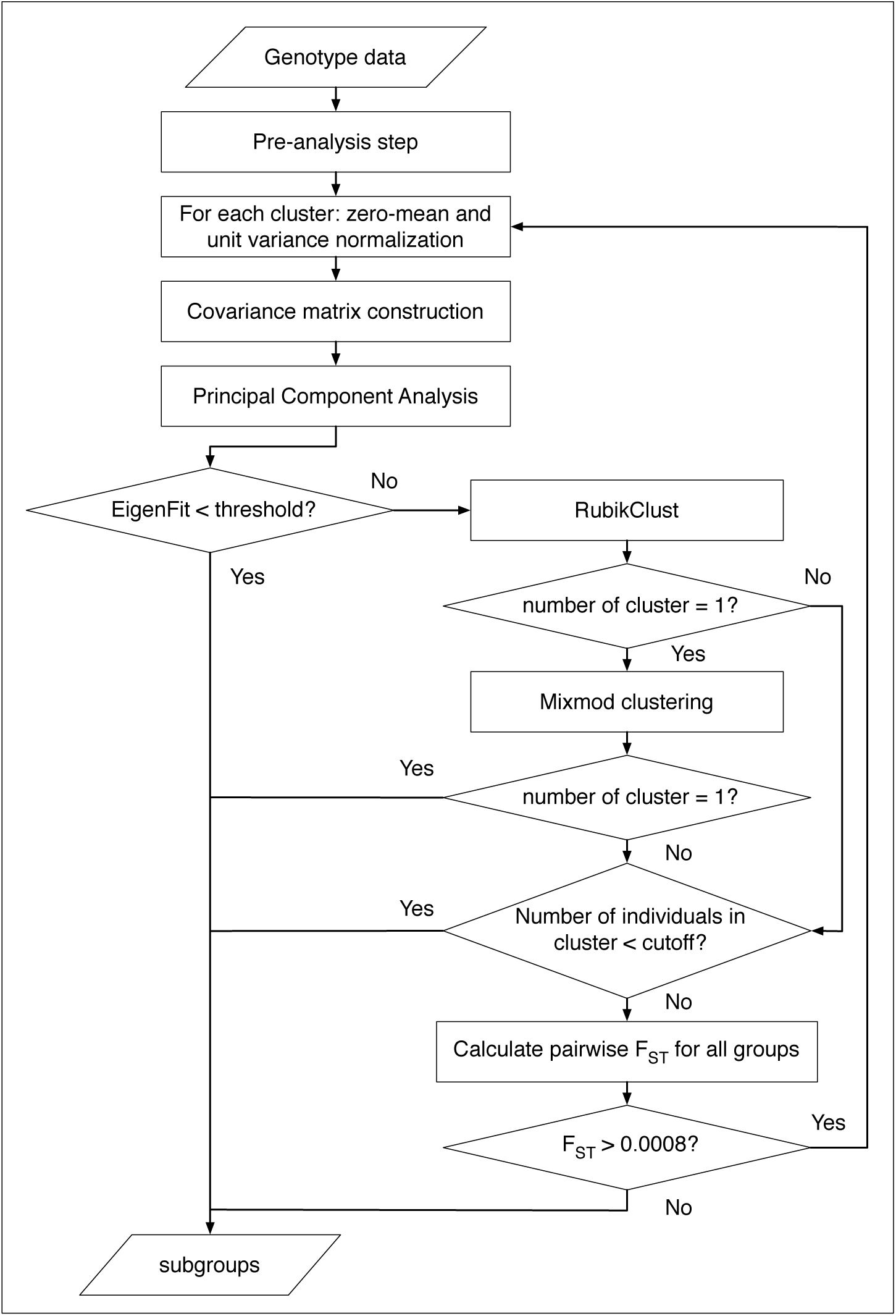
Flowchart for IPCAPS. The groups having individuals less than a cutoff, they are defined as outlying groups.

### Step 1

In each iteration, select genotype data *X* according to the remaining individuals from the previous iteration; however, a whole data are used for the first iteration. The matrix *X* contains *N* rows and *M* columns representing a number of individuals and a number of SNPs respectively. The SNP matrix is normalized using zero-mean and unit variance methods.

### Step 2

Construct a covariance matrix from matrix multiplication *XX^T^* in order to reduce to complexity for computation.

### Step 3

Extract principal components (PCs) from the matrix *XX^T^* as *XX^T^* = *UDU^T^*, where *U* represents eigenvectors and *D* is a diagonal matrix of positive eigenvalues of *XX^T^*of eigenvalues [2]. A matrix of eigenvectors is used as PCs. For faster computation, partial PCs can be obtained by the function *svds* from the R package *rARPACK* [11]. The matrix PCs contains *N* rows and *P* columns representing a number of individuals and a number of PCs respectively (see more details about *P* in the section of EigenFit).

### Step 4

Calculate the EigenFit value from the matrix *D* as described in the section of EigenFit. If the EigenFit is equal or less than a threshold, then stop the iteration and define the current set of individuals as a subgroup. If not, continue to the next step.

### Step 5

Apply RubikClust on derived PCs.

### Step 6

Check if the number of subgroups obtained from RubikClust equals 1. If so, then skip to step 9. If not, submit PCs to Mixmod clustering in step 7.

### Step 7

Apply to Mixmod clustering on PCs.

### Step 8

Check if the number of subgroups obtained from Mixmod clustering equals 1. If so, then stop this iteration and define the current set of individuals as a subgroup. If not, continue to the next step.

### Step 9

Check if the number of individuals in obtained subgroups is less than a cutoff. If so, then stop this iteration and define the current set of individuals as a subgroup. If not, continue to the next step.

### Step 10

Calculate pairwise F_ST_ for all pairs of subgroups. If F_ST_ is more than 0.0008, then continue to the next iteration in step 1. If not, the pairs with F_ST_ less than 0.0008 are combined and defined as a single (sub)group.

### Choices of internal clustering methods

IPCAPS relies on clustering individuals in an iterative way. There are several clustering methods exist. Cluster analysis can be achieved by various algorithms in different techniques depending on the notion of what constitutes clusters and how to efficiently detect them. Popular notions of clusters include dense areas of the data space, groups with small distances among members, and particular statistical distributions. Our objective is to detect fine-scale clusters based on multivariate estimation. In order to make a selection of clustering method to be used as integral part of the proposed IPCAPS methodology, we considered using the mouse-like dataset to examine clustering methods [12]. We selected unsupervised clustering algorithms to test with the mouse-like dataset; these algorithms include Mixmod [13], K-means, APCLUST [14], Mean shift [15], CLARA [16], PAM [17], hierarchical cluster analysis (HCA), and DBSCAN [18]. Since we aimed to select the most reasonable and fast-speed clustering method to integrate into our methodology, we submitted the mouse-like dataset to all methods for 100 times using the 64-bit OSX computer with the 1.3 GHz Intel Core i5 processor and 4 GB of memory. Finally, the average execution time of all methods was measured.

### Outlier detection: RubickClust

The quality of PCA-based clustering really depends on input data; hence, quality control process is needed at the beginning. Outliers are one factor that may disturb the quality of clustering result. Outliers can be detected using external tools as a part of QC process, for example using Rapid Outlier Detection [19]. However, we want to develop a method to eliminate outliers to improve the quality of IPCAPS result. We develop internal clustering method called RubikClust to detect rough structure, which aims to separate outliers into their own groups. The RubikClust algorithm is designed to apply to PCs hence PC1 is the most informative components, following by PC2 and PC3 consecutively. The RubikClust algorithm uses the concept of 3-dimension rotation to search for clear separation in all dimensions. The rotation is done by fixing PC3 axis (PC1 and PC2 are rotated), PC2 axis, and PC1 axis consecutively. The algorithm can be described as:

#### Step 1

Let *X_i_* and *Y_i_* be numeric vectors, where *i* = 1,2,3,…, *n* and *n* equals a number of samples. Calculate 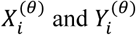, which are the rotated values of *X_i_* and *Y_i_* with angle *θ* where *θ* = 0,1,2,…,89, by using Equation (1) and (2) respectively.

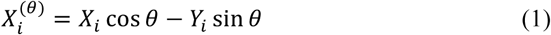

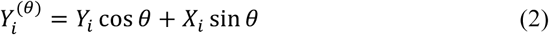

#### Step 2

Let 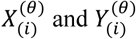 be the *i*-th sorted values.

#### Step 3

Normalize the sorted vector 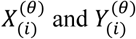 using Equation (3) and (4) respectively, then the normalized vector 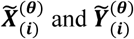 are obtained.

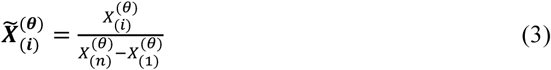

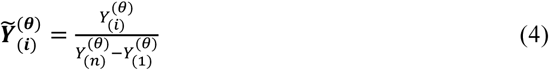

#### Step 4

Calculate the distance 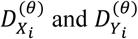 between two consecutive values of 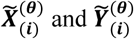 using Equation (5) and (6) respectively.

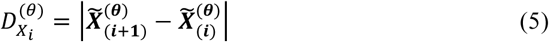

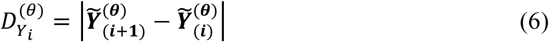

#### Step 5

Calculate the distance 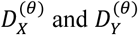 using Equation (7) and (9) respectively. Let *δ* be a pre-specified minimum distance.

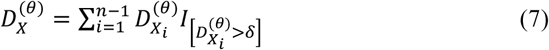

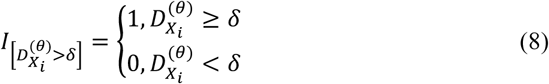

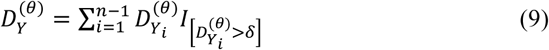

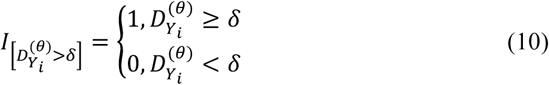

#### Step 6

Calculate 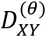 using Equation (11) for 0 - 89 degrees.

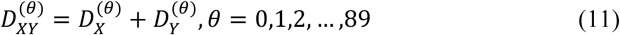

#### Step 7

Find the maximum value among 0 - 89 degrees and define as *D_max_*.

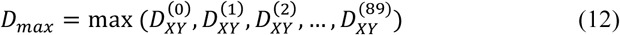

#### Step 8

Select the appropriate degree *θ* that corresponds to *D_max_,* let’s define it as the optimal degree *θ_opt_*.

#### Step 9

By fixing PC3, calculated the rotated PC1 and PC2 using Equation (13) and (14) respectively.

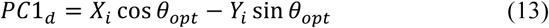

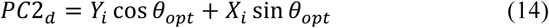

#### Step 10

Consecutively, fix the PC2 axis and PC1 axis, repeat step 1-9.

In Figure 3, we illustrate the 3D rotation method, which is applied in RubikClust. Unlike 3D visually rendering, the rotation in RubikClust is only applied for 0 - 89 degrees for clustering purpose (Figure 3 A). To determine clusters in each pair of PCs, *δ* is used as a cutoff for a minimum distance among consecutive projected values on each axis (Figure 3 B).

**Figure 3.**
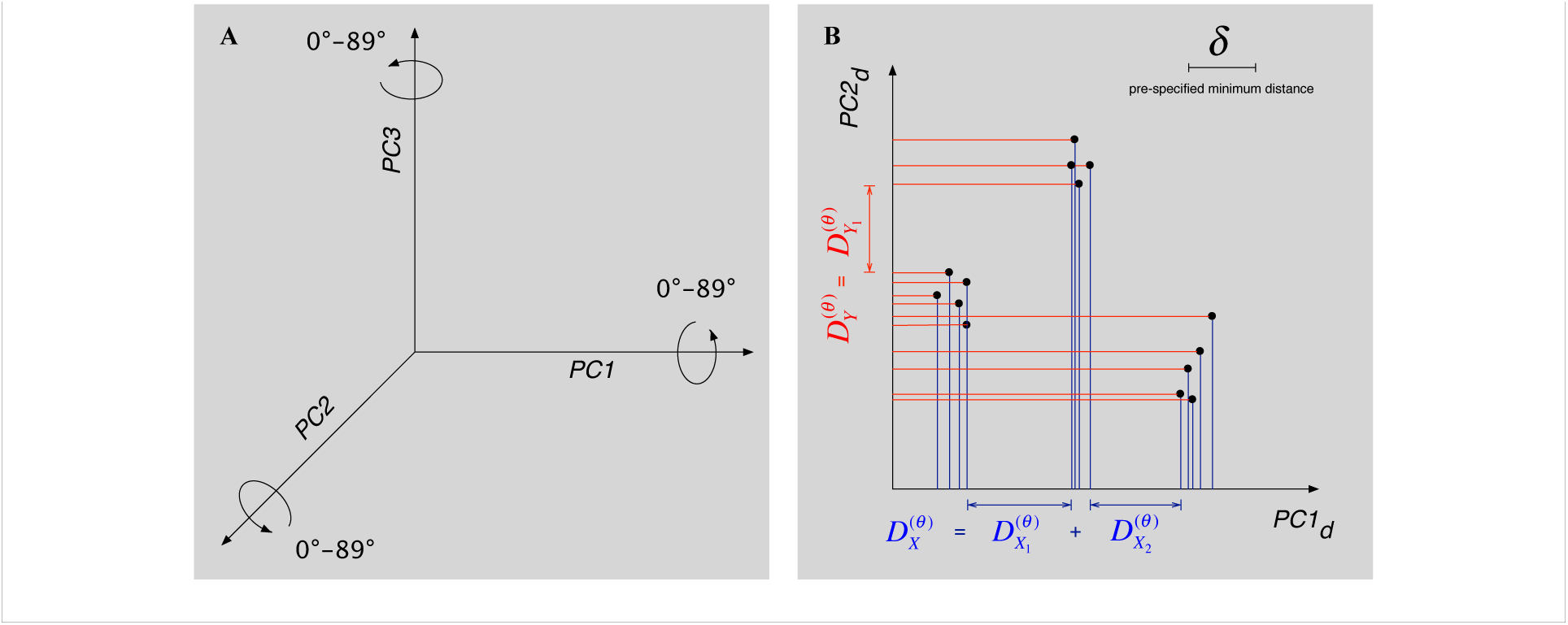
A: 3D rotation applied in RubikClust. B: In each rotation, here is a given example of distance determination for transformed PC1 and PC2.

We used the mouse-like dataset as reported in [12], and randomly added 7 outliers as well as a set of 20 individuals gathering outside the area of the mouse-like dataset. Then, we tested the ability of RubikClust for outlier detection with this dataset comparing to the other potential clustering methods such as APCLUST [14] and DBSCAN [18]. Lastly, we checked whether outliers could be separated from the mouse-like data. In addition, we wanted to compare the speed of these methods; therefore we performed 100 times of these experiments on the 64-bit OSX computer with the 1.3 GHz Intel Core i5 processor and 4 GB of memory. Finally, we measured the average execution time of the methods.

### Stopping criteria

Originally, the ipPCA methodology uses the *TW* statistic as a stopping criterion in the first version [6] and changes to use the EigenDev value in the second version [7]. Although ipPCA can capture general population structure, it cannot detect fine-scale structure when F_ST_ is closed to/or below 0.001 [20] such as between Swedish and Norwegian samples (F_ST_=0.001) or Polish and German samples (F_ST_=0.0012) [21].

Aiming for fine-scale structure detection, the ICAPS methodology combines 3 stopping criteria: 1) checking whether the EigenFit is lower than a threshold, 2) determining whether the F_ST_ value between two clusters is lower than a threshold, 3) checking whether a number of individuals in each cluster is lower than a cutoff. Note that the minimum value of cutoff is 3 according to the number of PCs used for RubikClust. If a minimum number of members is low (i.e. 3-5), then there is a chance to obtain many clusters with few individuals in each cluster. A minimum number of members can be defined arbitrarily, but it should not be too large (i.e. >20), depending on purposes whether small or large clusters are required to observe. When analyzing dataset with a small number of individuals, for example <100 individuals, it is more practical to consider a cutoff to a value between 5 to 10 individuals. Moreover, the IPCAPS methodology also relies on the Bayesian information criterion (BIC) [22] as in the function *mixmodCluster* in the R package *Rmixmod,* while achieving the optimal number of clusters [13].

### EigenFit

In IPCAPS, we propose a new stopping criterion called EigenFit, which is a maximum space between logarithms of eigenvalues. The EigenFit is defined by

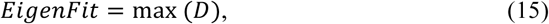

where max (*x*) is a function to obtain the maximum value of vector *x, D* is a vector of differences and is defined by

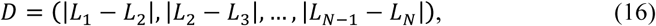

where *L_i_* is a vector of the logarithm of *N* eigenvalues or *L* = log (*eigenvalues*). Moreover, we define *P* as a proper number of PCs, which can be submitted to clustering method. Therefore, *P* is defined by

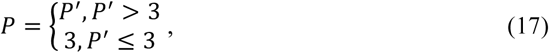

where *P′* is a estimated number of high impact PCs. However, if *P′* is lower than 3, PC1-3 are used for clustering. Here, *P′* is defined by

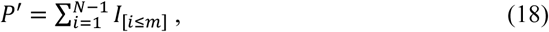

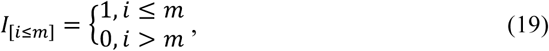

where *i* = 1,2,3, …, *N*, and *m* is an order of EigenFit in vector *D.* As defined in Equation (17), (18), and (19), the *P* first numbers of PCs are used in clustering step in order to improve speed for our algorithm. However, the minimum number of PCs to be submitted to clustering function is 3. The minimum threshold for EigenFit is motivated as 0.03, which can be observed from the results of simulated datasets as in the result section of simulation scenario II. The maximum threshold for EigenFit is motivated as 0.18, which can be observed from the HapMap populations.

### Optimal Fixation Index

The fixation index (F_ST_) can be used to measure a distance between populations. Although F_ST_ = 0.001 was referred as the genetic distance between close populations within Europe [21,23], it is still unclear how to quantify fine-scale structure. Here, we set up the experiments to check the lowest F_ST_ that can be related to fine-scale structure (different from no structure at all).

For visualization purposes, we firstly calculated PCs from 6 simulated datasets of 2 populations with F_ST_ values were set as 0.001, 0.002, 0.003, 0.004, 0.005, and 0.006. These datasets contain 10,000 SNPs and 500 individuals (250 individuals for each population). We used FilestSim to simulate the data.

Secondly, we simulated 100 replicates of one population with 5,000, 10,000, and 20,000 SNPs, and 500, 1,000, 2,000, 4,000, 6,000, and 8,000 individuals (i.e., 3x6 settings). We forced the data to be separated into two groups using K-means clustering. Later, we calculated F_ST_ for these two clusters, and then checked for the minimum, the maximum, and the average. In particular, Hudson’s method was used to estimate F_ST_ because this method is more stable than the methods of Nei, and Weir and Cockerham [24]. The method does not overestimate F_ST_ for unbalanced sample sizes. The equation for Hudson’s method was originally given by

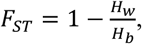

where *H_w_* is the mean number of differences within populations, and *H_b_* is the mean number of differences between populations. Hudson did not give explicit equations for *H_w_* and *H_b_*. Therefore, Bhatia derived the following equation [24] for F_ST_.

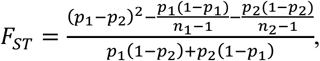

where *n_i_* is the sample size and *p_i_* is the sample allele frequency in population *i* for *i* ∈ {1,2}. To estimate F_ST_ in our experiments we used the formulation in Equation (2.5.2.2).

### Missing data handling

Missing genotypes can be treated differently, for example, by removal from the data [4], by ignorance [6], by common value substitution [10], by (multiple) imputation [25]. The validity of these strategies depends on the underlying process of missing data generation. Three such processes exist: missing completely at random (MCAR), missing at random (MAR) or non-random missingness (MNAR). Whatever the missing data process is, during the quality control process, it is best to keep missingness rates to low levels. In human genetic association analyses, we most commonly do so by using imputation strategies. These may or may not rely on reference data such as the HapMap [26] or data from the 1000 Genomes Project [27]. Several statistical methods for imputing genotypes exist [28], including KNN imputation [29] and imputation via prediction models [30,31]. In this thesis, and to reduce computational complexity, we consistently imputed missing genotypes by their most common value, unless stated otherwise.

Note that in the context of population structure detection, where the aim is to highlight fine-scale structure, it is less clear which reference data for imputation to use. Formulating recommendations regarding this are needed but are beyond the scope of this thesis. As single value imputation is only valid under MCAR processes, all of our practical data analyses were supplemented by a basic missing data process analysis.

### Performance assessment using different simulation scenarios

To test the performance of IPCAPS, we set up several experiments according to 4 different scenarios; simulation scenario I aims to investigate type I error, simulation scenario II aims to assess the accuracy of IPCAPS, simulation scenario III targets quantifying scalability and speed, and simulation scenario IV allows to investigate IPCAPS capability for complex data structures. To generate the required synthetic data, we have built our own in-house simulator, called FilestSim. The cluster agreement between two clustering results is assessed via the Adjusted Rand Index (ARI). The ARI function is available in the R Package *mclust* [32]. The highest ARI value is 1, which represents identical matching between 2 groups, while negative value represents mismatch.

### Simulation scenario I: test for type I error

The objective of simulation scenario I is to examine the type I error rate of our method. Ideally, there shouldn't be any error if we apply IPCAPS on a single homogeneous population. In other words, in this case, the IPCAPS algorithm should only reveal 1 group; the initial population. We compared our method to other iterative pruning based clustering methods such as ipPCA [6,7], iNJclust [32], and SHIPS [33]. Moreover, we simulated 1 population with 500 individuals and 10,000 SNPs without any outliers and did so 100 times (i.e., 100 replicates). The parameter settings for FilestSim are listed in Table 1.

**Table 1.** Parameter settings for simulation scenario I

| Parameters | Settings |
| --- | --- |
| Number of populations | 1 |
| Number of individuals | 500 |
| Number of SNPs | 10,000 |
| Number of outliers | 0 |
| Number of replicates | 100 |

### Simula tion scenario II: test for a ccura cy

The objective of simulation scenario II is to determine the accuracy of IPCAPS. Comparative iterative pruning based methods for clustering are the same as for simulation scenario I: ipPCA, iNJclust, and SHIPS. For scenario II we simulated 100 replicates of 10,000 SNPs and 500 individuals per population. For the settings SII-1, SII-3 and SII-5, 2 populations were simulated. We added an additional population in the settings SII-2, SII-4 and SII-6. The adopted F_ST_ values represent pairwise genetic distances as before (Hudson’s fixation index) and ranged from 0.0008 to 0.005. We selected the lowest F_ST_ as 0.0008 according to the result of simulation s, and the highest F_ST_ as 0.005 according to the genetic distance among clearly distinct European populations [21,23]. To assess the impact of outliers, we added 3 outliers in the settings SII-3 and SII-4, and 5 outliers in the settings SII-5 and SII-6. Particularly, an outlier is considered that it has been detected when it is separated into its own groups or is grouped with other outliers. All setting parameters are summarized in Table 2.

**Table 2.** Parameter settings for simulation scenario II

| Parameters | Settings |  |  |  |  |  |
| --- | --- | --- | --- | --- | --- | --- |
|  | SII-1 | SII-2 | SII-3 | SII-4 | SII-5 | SII-6 |
| Number of populations | 2 | 3 | 2 | 3 | 2 | 3 |
| Distance ($F_{ST}$) between populations | 0.0008, 0.0009, 0.001, 0.002, 0.003, 0.004, 0.005 | | | | | |
| Number of individuals per population | 500 |  |  |  |  |  |
| Number of SNPs | 10,000 |  |  |  |  |  |
| Number of outliers | 0 | 0 | 3 | 3 | 5 | 5 |
| Number of replicates | 100 |  |  |  |  |  |

### Simula tion scenario III: test for scalability and speed

The objective of scenario III is to check the scalability and speed of IPCAPS. In particular, we want to investigate which of the two, the number of individuals or the number of SNPs, has the most impact on computation time. According to the results of scenario II, we chose to compare IPCAPS to ipPCA only. The parameter settings for this simulation scenario are as follows. We simulated 100 replicates of 2 populations while fixing F_ST_ at 0.005. This single fixed value of F_ST_ is motivated by the fact that IPCAPS is able to accurately separate 2 populations with F_ST_=0.005 (see scenario II.) For setting SIII-1, we fixed the number of input SNP to 10,000 and varied the number of individuals from 100 to 10,000. For setting SIII-2, we considered 1,000 individuals and varied the number of SNPs from 25,000 to 100,000. To measure the performance of IPCAS in terms of computation time, we performed all experiments on the same 64-bit Linux cluster with 2.2 GHz Intel Xeon 8-core processor and 128 GB of memory per node. Since the cluster was working routinely and we couldn’t control other running processes, we reported the median of execution times from 100 replicates instead of the mean. All parameter settings are summarized in Table 3.

**Table 3.** Parameter settings for simulation scenario III

| Parameters | Settings |  |
| --- | --- | --- |
|  | SIII-1 | SIII-2 |
| Number of populations | 2 |  |
| Distance ($F_{ST}$) between populations | 0.005 | |
| Number of individuals (2 populations are balanced) | 100, 2,500, 5,000, 7,500, 10,000 | 1,000 |
| Number of SNPs | 10,000 | 25,000, 50,000, 75,000, 100,000 |
| Number of replicates | 100 |  |

### Simulation scenario IV: test for dealing with increased data complexity

In real-life applications, data are expected to be complex and exhibit peculiar or complex substructures, as well as outlying individuals. The objective of simulation scenario IV is to test the capability of our method in complex data structures. To this end we simulated 10,000 SNPs and 2,400 individuals consisting of 8 populations (300 individuals each). We also randomly added 20 outliers. All parameter settings are shown in Table 4. The estimated pairwise F_ST_ values obtained from FilestSim are listed in Table 5.

**Table 4.** Parameters to FilestSim for simulation scenario IV

| Parameters | Settings |
| --- | --- |
| Number of populations | 8 |
| Number of individuals | 2,400 (300 individuals each) |
| Number of outliers | 20 |
| Number of SNPs | 10,000 |
| Distance from the first population ($F_{ST}$) | pop2 (0.002), pop3 (0.004), pop5 (0.006), pop6 (0.05), pop8 (0.05), pop9 (0.01), pop10 (0.008) |

**Table 5.** Estimated F_ST_ of simulated data IV

|  |  |  |  |  |  |  |  |
| --- | --- | --- | --- | --- | --- | --- | --- |
| pop2 | 0.0021 |  |  |  |  |  |  |
| pop3 | 0.0040 | 0.0042 |  |  |  |  |  |
| pop5 | 0.0062 | 0.0062 | 0.0081 |  |  |  |  |
| pop6 | 0.0499 | 0.0502 | 0.0526 | 0.0550 |  |  |  |
| pop8 | 0.0505 | 0.0506 | 0.0533 | 0.0541 | 0.1005 |  |  |
| pop9 | 0.0101 | 0.0100 | 0.0124 | 0.0144 | 0.0595 | 0.0591 |  |
| pop10 | 0.0084 | 0.0081 | 0.0104 | 0.0127 | 0.0580 | 0.0551 | 0.0160 |
|  | pop1 | pop2 | pop3 | pop5 | pop6 | pop8 | pop9 |

## RESULTS

This section reports all main results related to the development of IPCAPS and its performance.

### Comparative study on cl ustering methods

Using the mouse-like dataset, most methods can detect 3 clusters (blue, red, and green), except DBSCAN, which is able to only detect 1 cluster (Figure 4). The Mixmod algorithm provides the most realistic results compared to the other methods. Although K-means, APCLUST, Mean shift, CLARA, PAM, and HCA are able to identify 3 clusters, the results are not well separated and it is difficult to observe clusters’ boundary without colors. Regarding computation time, most algorithms only took a few seconds (<5 sec) to carry out the clustering, except for the Mean shift and APCLUST algorithms, which took more than 100 seconds.

**Figure 4.**
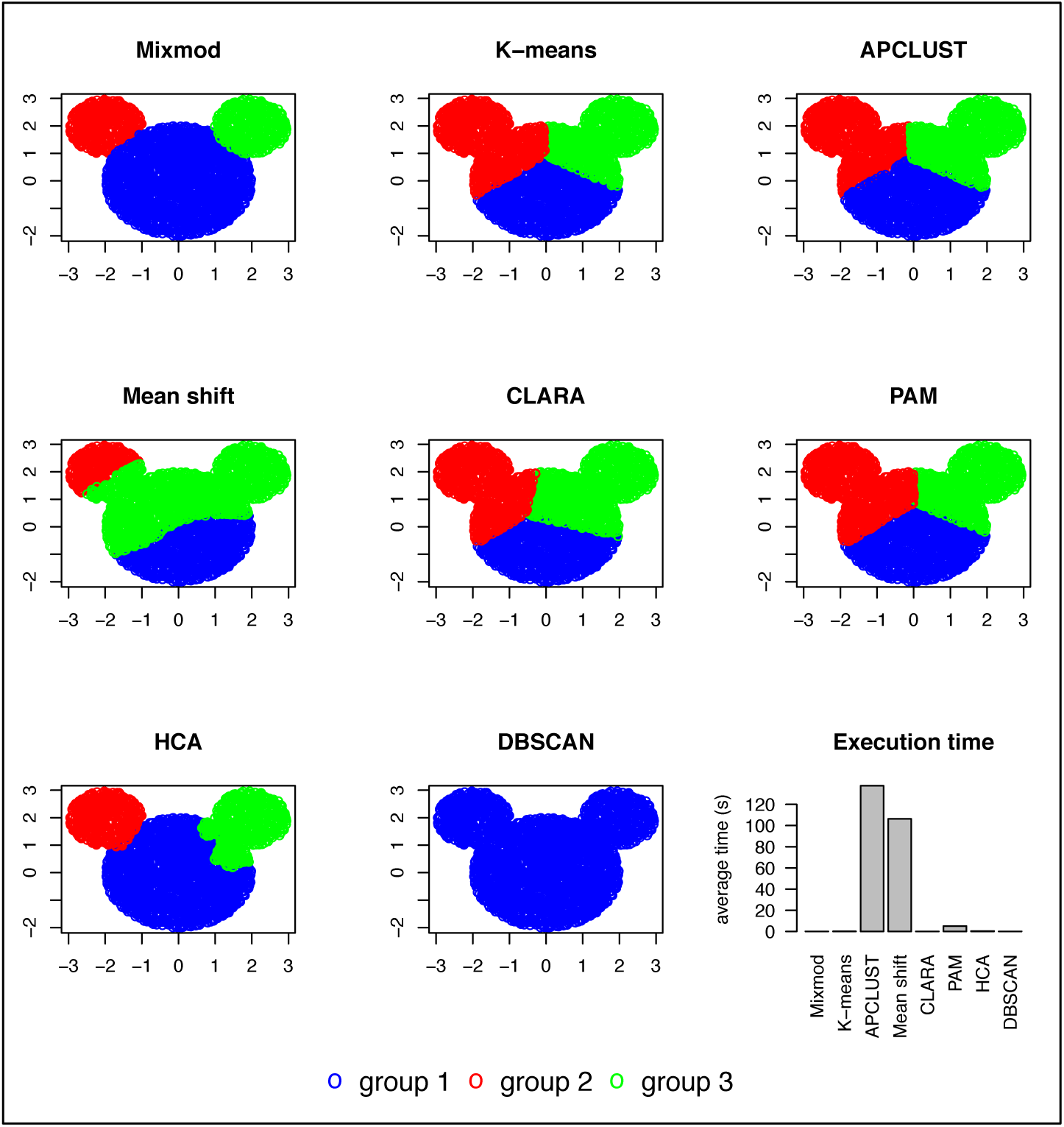
Results on the mouse-like dataset for clustering algorithms.

### Comparative study on outlier detection

As can be seen from Figure 5, RubikClust and DBSCAN can accurately detect all additional data added to the mouse-like dataset. Moreover, RubikClust can also detect the added group 8 (Figure 5, brown), while DBSCAN cannot separate this group from other outliers. The APCLUST algorithm poorly performs on the as it is unable to discriminate the outliers from the mouse-like dataset. Regarding the required computation time, RubikClust and DBSCAN need less than 10 seconds. This is much faster than APCLUST, which needs more than 160 seconds.

**Figure 5.**
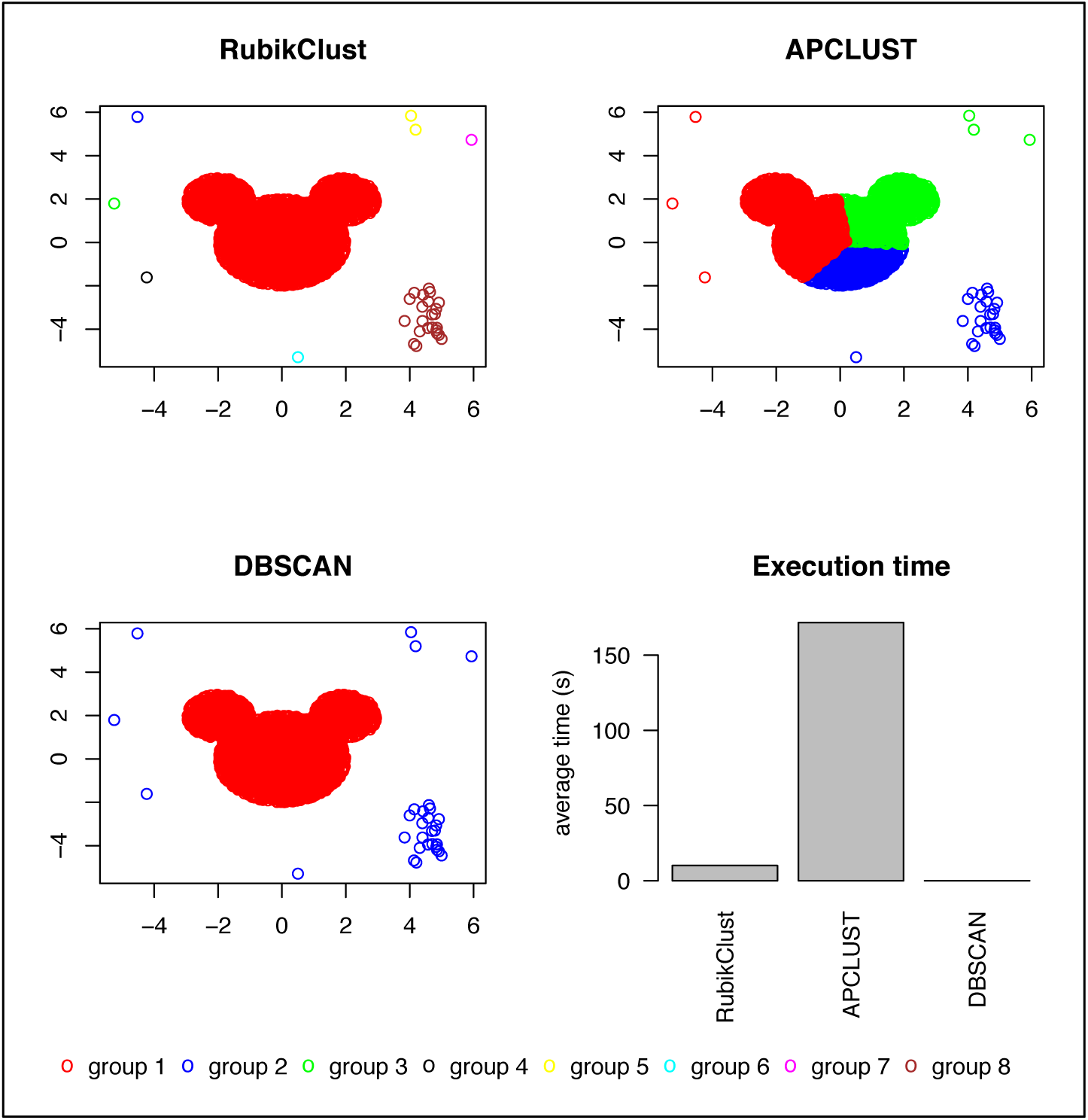
Results on the mouse-like dataset and simulated outliers.

### Threshold for fine-scale structure

As explained in the section of stopping criteria, the considered F_ST_ values resemble observable F_ST_ values for European subpopulations [21,23]. As can be seen from Figure 6, the first two PC components are unable to distinguish between pop1 (red) and pop2 (blue) when F_ST_=0.001. This setting mimics two closely related populations, such as the Swedish (SW) and the Norwegians (NO). In the case of F_ST_=0.002, pop1 and pop2 are not clearly separated either and are hardly separable without coloring. This is like the close populations such as Swedish and Hungarian (HU). In the case of F_ST_=0.003, the separation is observable between pop1 and pop2 via PC1-2. This setting resembles distinguishing between Spanish (SP) and Hungarian European subpopulations. For F_ST_>=0.004, pop1 and pop2 are separated, as Swedish vs Romanian, Spanish vs Polish (PO), and Spanish vs Russian (RU) in Figure 6.

**Figure 6.**
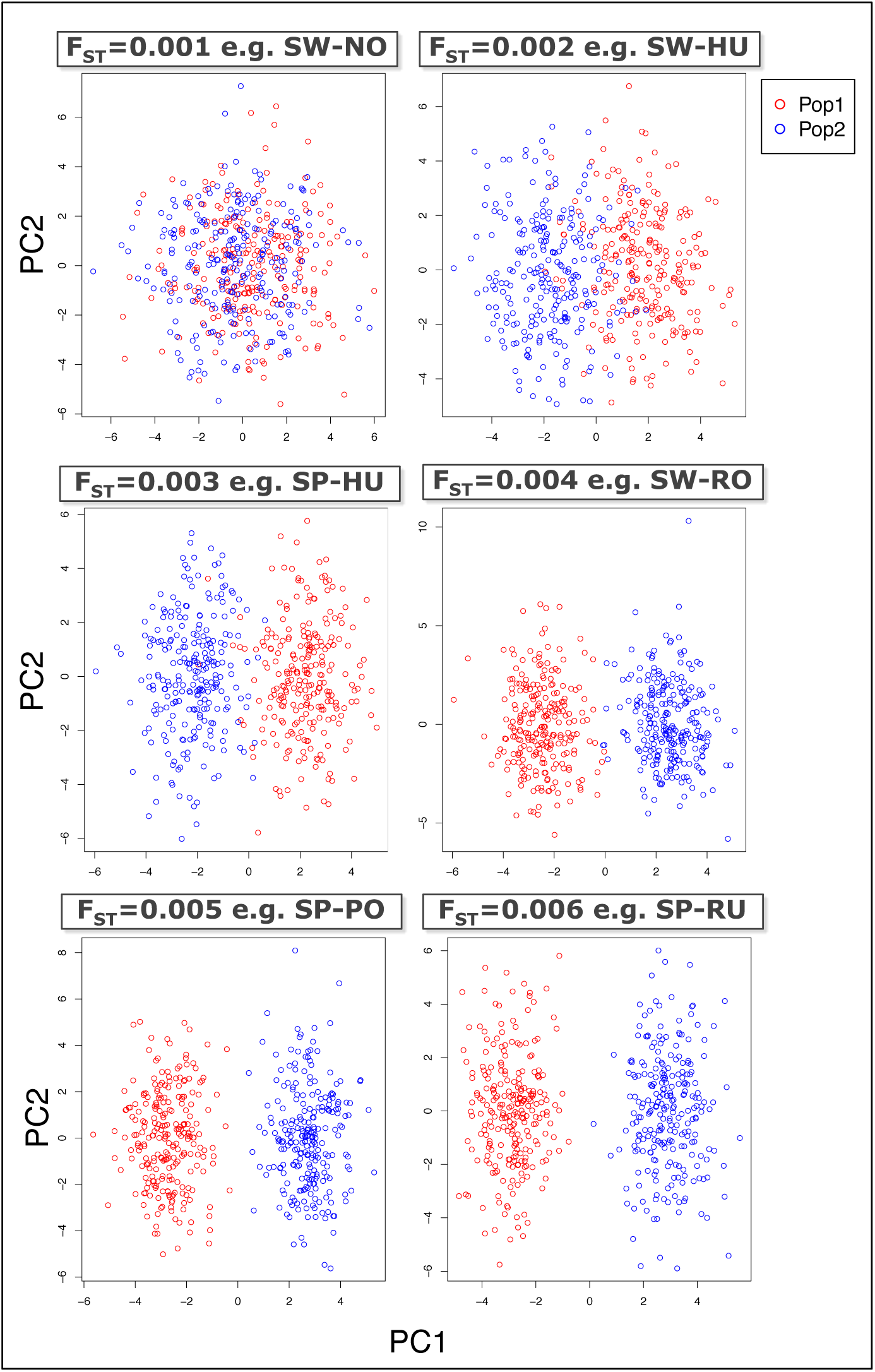
Visualization of PC1-2 on simulated data with different F_ST_ values.

Furthermore, Figure 7 shows the estimated F_ST_ values according to sample size (i.e., number of individuals in a single population) and by different numbers of SNPs (5K, 10K and 20K). The black curves follow the average estimated F_ST_ over 100 replicates, for a particular combination of sample size and number of SNPs. The blue areas are determined by the minimum and maximum estimated F_ST_ corresponding to an average F_ST_ above. Overall, we observe that the average F_ST_ trends to decrease when the number of individuals and the number of SNPs increases. The highest F_ST_ among all experiments is around 0.0008. This overestimation for the expected F_ST_ of zero was obtained for the smallest dataset, involving 5,000 SNPs and 500 individuals.

**Figure 7.**
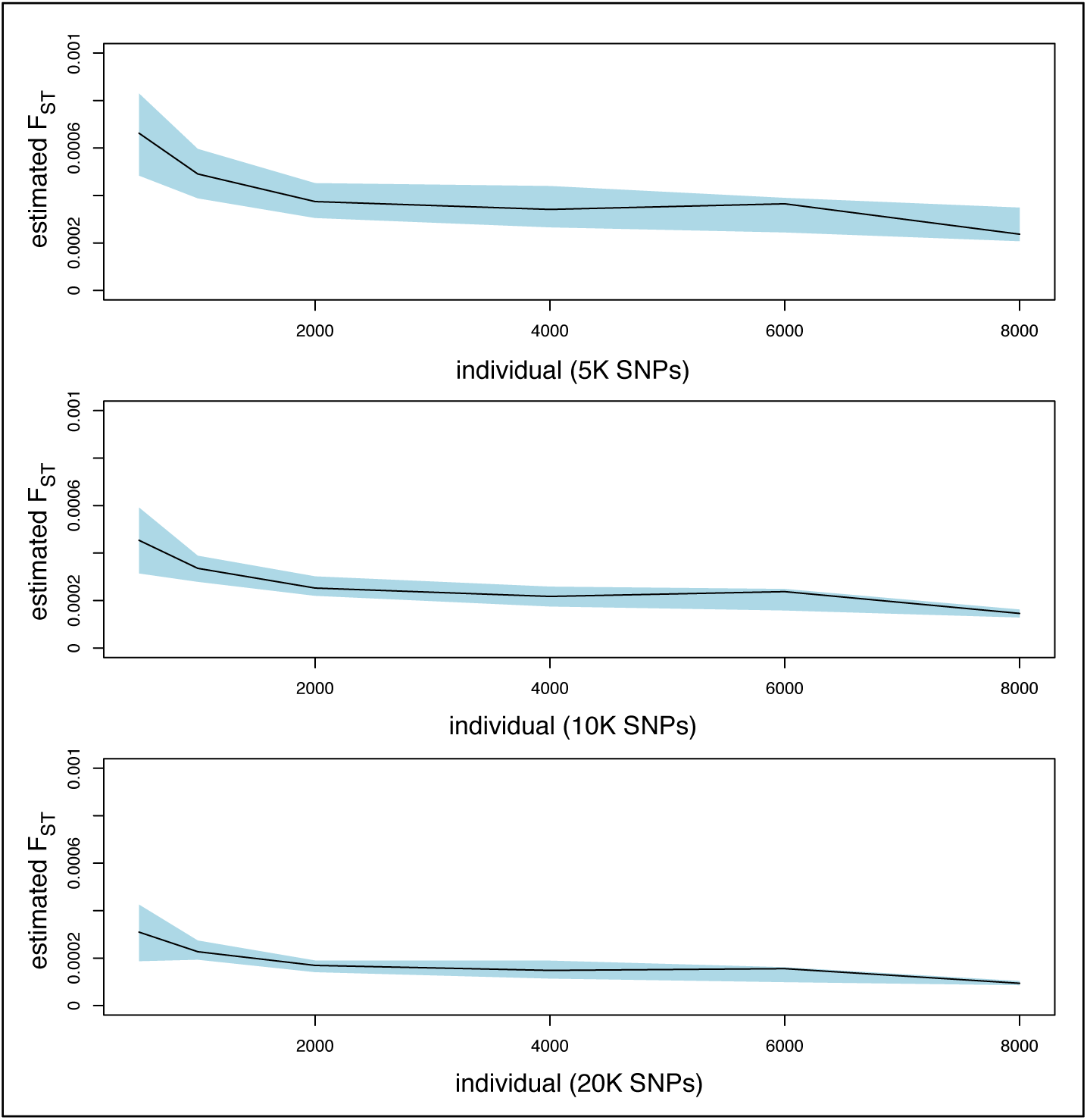
Result of experiment for determining a threshold for fine-scale structure. The black lines are the average of estimated F_ST_ and the blue areas are the area between the minimum and maximum of estimated F_ST_.

### Type I error (simulation scenario I)

From the considered clustering methods, as explained in the simulation scenario I, only IPCAPS and SHIPS do not split up the single population in subgroups (average ARI=1). Notably, ipPCA and iNJclust enforce 2 subgroups (average ARI=0) and 172.79 groups (average ARI=0), respectively.

**Table 6.** Clustering result of experiments on type I error

| Methods | Average ARI | Average number of clusters |
| --- | --- | --- |
| IPCAPS | 1.00 | 1.00 |
| ipPCA | 0.00 | 2.00 |
| iNJclust | 0.00 | 174.79 |
| SHIPS | 1.00 | 1.00 |

### Accuracy (simulation scenario II)

Overall, IPCAPS (red curve in Figure 8) has optimal performance compared to ipPCA (blue), SHIPS (green) and iNJclust (yellow), in terms of accuracy expressed by average ARI estimated over 100 replicates when comparing observed and expected clustering methods. In particular, IPCAPS performs well with 100% accuracy when F_ST_=0.002, for all simulation scenarios, while the other strategies perform less for the same F_ST_=0.002. As for the other strategies, the performance of IPCAPS decreases for decreasing F_ST_<0.002 although the accuracy reduction is least dramatic for IPCAPS compared to ipPCA, SHIPS and iNJclust.

**Figure 8.**
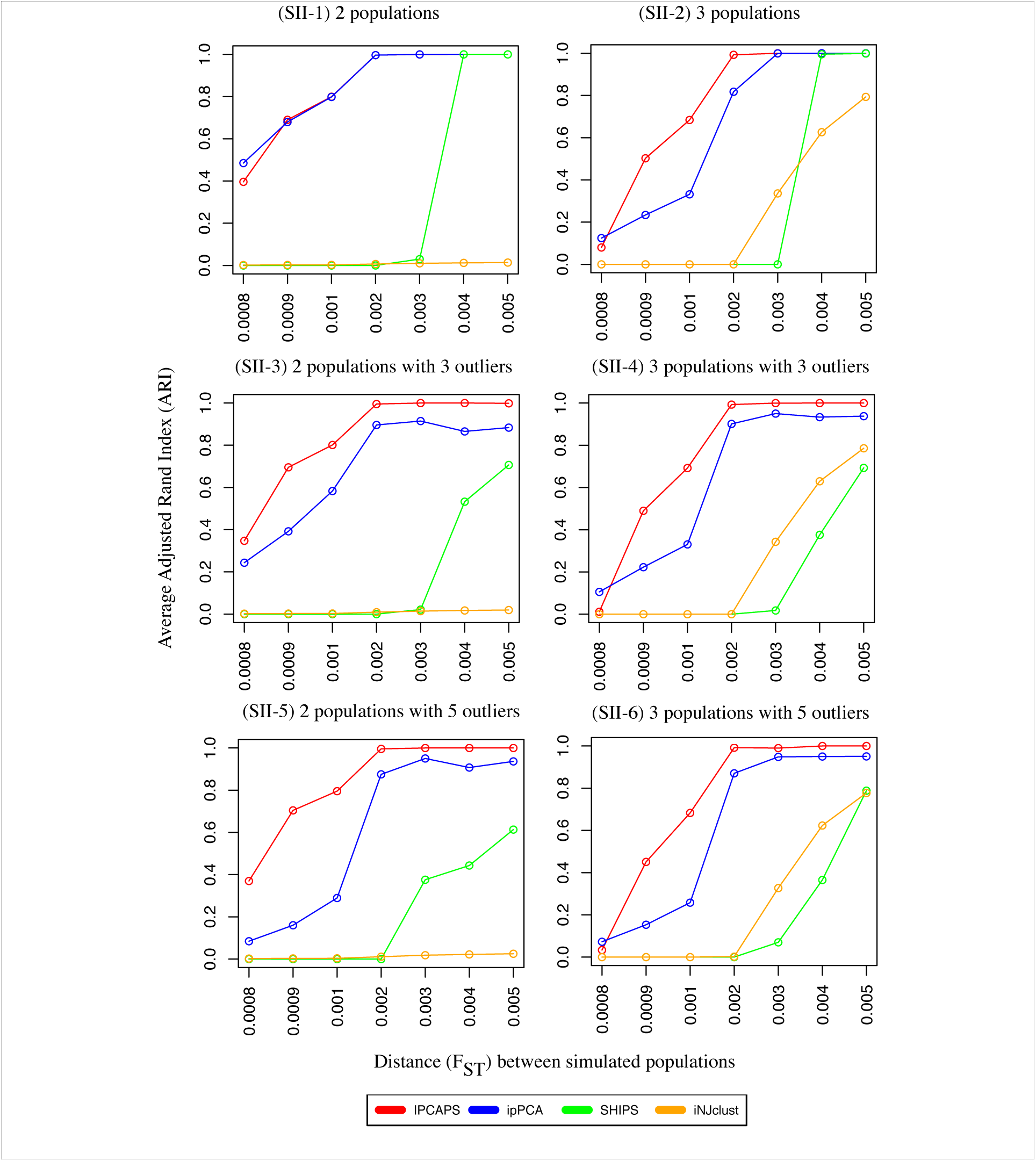
Accuracy of IPCAPS comparing to other tools.

In the case of 2 populations without outliers (setting SII-1), IPCAPS and ipPCA give similar result, but ipPCA performs slightly better for F_ST_=0.0008. The average ARI of both methods increases to 0.8 when F_ST_=0.001, and reaches 1 when F_ST_=0.002. SHIPS becomes highly accurate (ARI=1) only when F_ST_>=0.004, while iNJclust poorly performs (ARI=0) in this setting.

In the case of 3 populations without outliers (setting SII-2), IPCAPS is more accurate than the other considered methods. The average ARI of IPCAPS reaches 1 when F_ST_=0.002, while the performance of ipPCA drops in this setting with ARI reaching 1 from F_ST_=0.003 onwards. SHIPS shows similar performance in setting SII-2 as in the previous setting SII-1. The average ARI of iNJclust starts to increase when F_ST_=0.003 and increases up to 0.8 but never reaches 1.

In the case of 2 populations with 3 and 5 outliers (settings SII-3 and SII-5), IPCAPS maintains its good performance, similar to the simulation setting SII-1 with 2 populations and no outliers. The performances of ipPCA and SHIPS drop compared to setting SII-1; iNJclust consistently performs poorly for all F_ST_ in this setting (ARI=0).

In the case of 3 populations with 3 and 5 outliers (settings SII-4 and SII-6), IPCAPS still shows similar performance to the corresponding settings without outliers. The performances of ipPCA and SHIPS drop compared to setting SII-2. Interestingly, iNJclust shows increased accuracy for F_ST_>0.003 However, the average values of ARI of ipPCA, SHIPS and iNJclust always stay lower than 1 in settings SII-4 and SII-6.

Figure 9, focusing on simulation settings with outliers and visualizing the number of outliers detected versus F_ST_, clearly shows that IPCAPS is able to detect the largest number of outliers compared to ipPCA, SHIPS and iNJclust. Recall that an outlier is considered that it has been detected when it is separated into its own groups or is grouped with other outliers. Particularly in the settings SII-4 and SII-6, iNJclust has a hard time in identifying any outlier at all. Although ipPCA is able to identify outliers, for all scenarios, it can detect approximately 2 out of 3 in the settings SII-3 and SII-4, and 4 out of 5 in the settings SII-5 and SII-6. SHIPS cannot detect outliers in the settings SII-3 and SII-4, but it is able to identify approximately 1 out of 5 in the setting SII-5 and SII-6.

**Figure 9.**
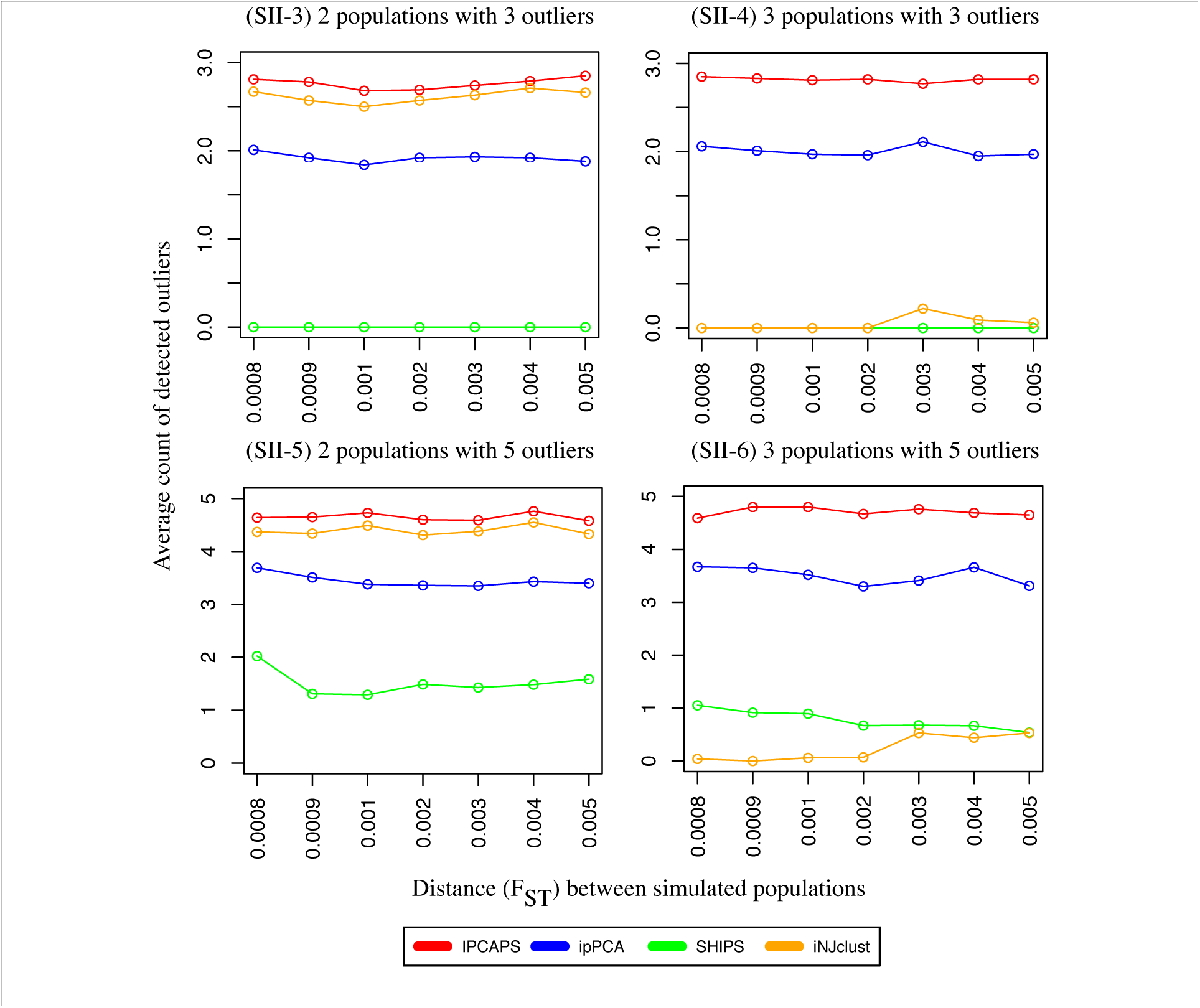
Performance on outlier detection.

### Salability and speed (simulation scenario III)

The average execution time of ipPCA (Figure 10. - blue curve) exponentially grows according to the number of individuals, reaching >24,000 seconds for 10,000 individuals (setting SIII-1). In contrast, the average execution time of IPCAPS (Figure 10. - red curve) is lower than ipPCA; it reaches approximately 2,000 seconds for 10,000 individuals (setting SIII-1). For setting SIII-2, the average execution time of IPCAPS and ipPCA is much lower than for setting SIII-1. The average execution time of ipPCA reaches 150 seconds for 100K SNPs, while the average execution time of IPCAPS is slightly lower and less than 150 seconds for 100K SNPs.

**Figure 10.**
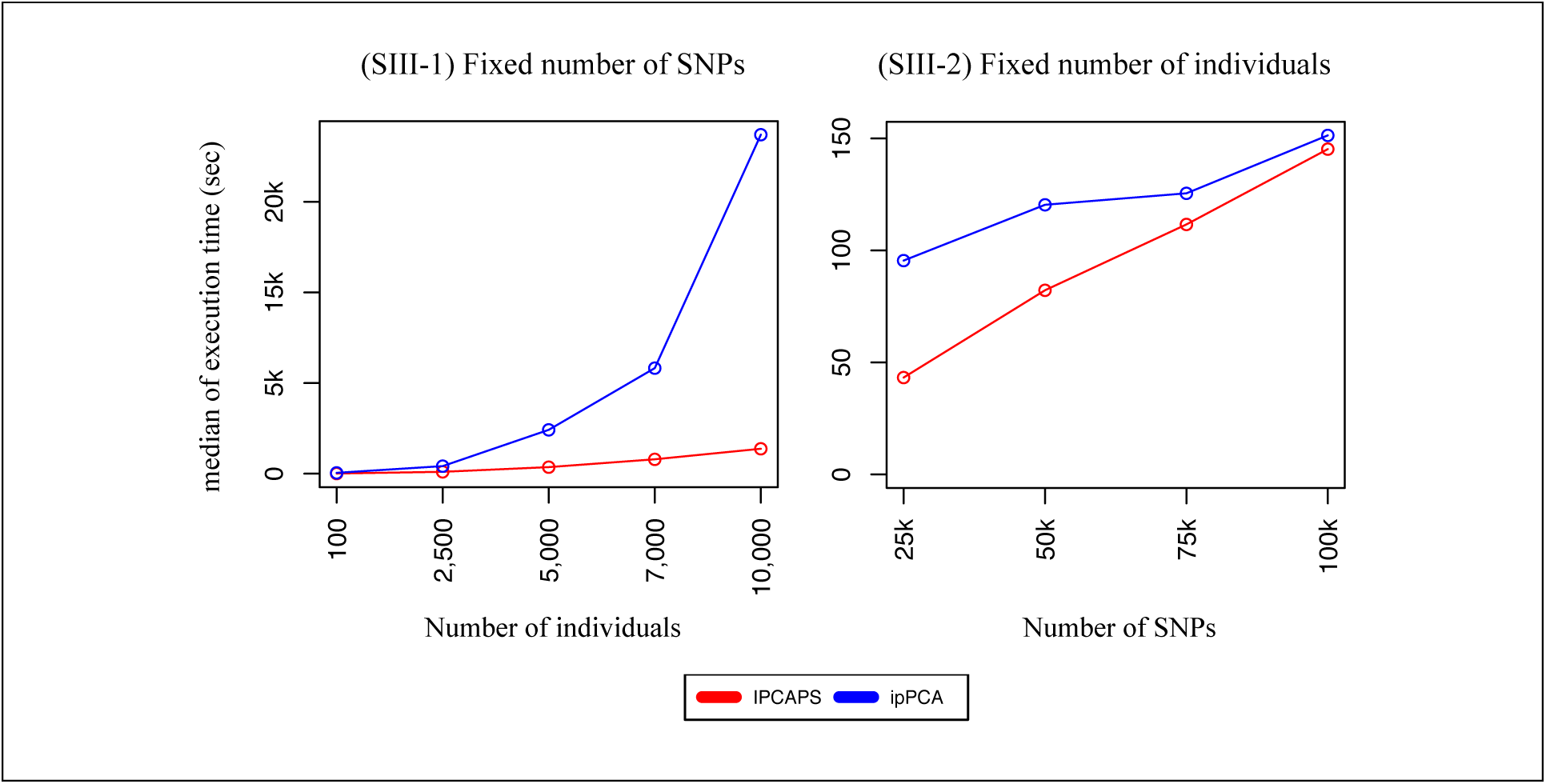
Execution time of IPCAPS and ipPCA.

### Increased da ta complexity (simula tion scenario III)

For this simulated dataset, IPCAPS can perform clustering with ARI=0.99, and clustering result is shown in Table 7. We observe that that pop1, pop2, pop3, pop5, pop6, pop8, pop9, and pop10 are mainly assigned to their own groups (group 1-8). The outliers 4 and 7 are occasionally assigned to groups 9 through 13.

**Table 7.** Clustering result of simulation scenario IV

| Groups | pop1 | pop10 | pop2 | pop3 | pop5 | pop6 | pop8 | pop9 | outlier4 | outlier7 |
| --- | --- | --- | --- | --- | --- | --- | --- | --- | --- | --- |
| 1 | 0 | 0 | 0 | 0 | 0 | 300 | 0 | 0 | 0 | 0 |
| 2 | 0 | 0 | 0 | 0 | 0 | 0 | 300 | 0 | 0 | 0 |
| 3 | 0 | 0 | 0 | 0 | 0 | 0 | 0 | 300 | 0 | 0 |
| 4 | 0 | 300 | 0 | 0 | 0 | 0 | 0 | 0 | 0 | 0 |
| 5 | 0 | 0 | 0 | 0 | 300 | 0 | 0 | 0 | 0 | 0 |
| 6 | 287 | 0 | 18 | 0 | 0 | 0 | 0 | 0 | 0 | 0 |
| 7 | 0 | 0 | 0 | 300 | 0 | 0 | 0 | 0 | 0 | 0 |
| 8 | 13 | 0 | 282 | 0 | 0 | 0 | 0 | 0 | 0 | 0 |
| 9 | 0 | 0 | 0 | 0 | 0 | 0 | 0 | 0 | 1 | 2 |
| 10 | 0 | 0 | 0 | 0 | 0 | 0 | 0 | 0 | 2 | 1 |
| 11 | 0 | 0 | 0 | 0 | 0 | 0 | 0 | 0 | 2 | 2 |
| 12 | 0 | 0 | 0 | 0 | 0 | 0 | 0 | 0 | 2 | 2 |
| 13 | 0 | 0 | 0 | 0 | 0 | 0 | 0 | 0 | 3 | 3 |

## DISCUSSION

Regarding the clustering methodology used in IPCAPS, we first considered clustering algorithms that were available in the R language. After testing, Mixmod was preferred because it performed well in complex datasets such as the mouse-like data [12]. Generally, the PCs representing individuals from the same population are packed, which can be observed from simulated data. We potentially see that Mixmod well suits our purpose. We furthermore developed an algorithm called RubikClust that enables the identification of outliers. These outliers are removed prior to the adoption of Mixmod, so as to increase the stability of IPCAPS results. Once dealing with other omics or multi-omics, these methods may need to be replaced for further improvement to obtain consistent clusters (e.g., Consensus Clustering [34], ConsensusClusterPlus [35], and COCA [36]).

For stopping criteria of IPCAPS’s iterative process, we combine 3 criteria to terminate the iterative process, determining the minimum number of individuals in a group, using the EigenFit heuristic, and using F_ST_. We confirmed that that F_ST_ estimated from large samples sizes is more precise, while from small samples sizes is overestimated. If there are a sufficient number of markers, F_ST_ is estimated precisely [37]. Although F_ST_=0 is referred as “no genetic differentiation”, it is clearly that F_ST_ depends on sample size and a number of markers. Thus, a threshold is needed to be refined and it should be more than zero. According to our experiments, we observe that F_ST_=0.0008 is an appropriate threshold to terminate the iterative process in IPCAPS.

Estimated Type I error rates for IPCAPS in our simulation scenario are zero due to the fact that IPCAPS has adopted the F_ST_ threshold according to samples size and a number of SNPs. Accuracy is defined as the ability to retrieve existing substructure, the adjusted rand index method (ARI) is used to measure accuracy [38]. In all simulation scenarios, IPCAPS generally outperforms ipPCA, SHIPS and iNJclust. The ipPCA method has higher average ARI than IPCAPS for F_ST_=0.008, because ipPCA over separates data as it is observed that the type I error of ipPCA is high (100%). When testing for speed, it is observed that IPCAPS is faster than ipPCA. When dealing with complex population structure in our simulation scenario, IPCAPS delivers satisfied results. In general, IPCAPS has excellent performance for both fine-scale and large-scale settings supporting from experimental results. Initially, the objective of this thesis is to target a tool that deals with fine-scale structure. IPCAPS accurately deal with the rough structures. Thus, it is necessity to split off big groups and then to zoom in for finding additional subtle structures.

Once dealing with outliers, data complexity increases. In general, an outlier is an observation that is isolated from other observations in PC space. IPCAPS uses RubikClust algorithm to detect outliers. Other efficient tools for outlier detection exist, such as Rapid Distance-Based Outlier Detection [19] and the Anselin Local Moran’s method [39]. The impact of outliers on clustering results with IPCAPS was assessed via several simulation studies, which entailed increasingly complex data structures. Using the mouse-like data with outliers, IPCAPS could clearly separate outliers and small group of points from the mouse-like data points. In simulation scenario with outliers for accuracy, IPCAPS could averagely detect outliers more accurate than ipPCA, iNJclust and SHIPS. Once dealing with increased complexity data, IPCAPS could detect outliers even though outliers couldn't be observed using PC1-2.

Lastly, the computational burden of IPCAPS depends on the number of individuals due to the dimensionality reduction prior to PCA via the *XX^T^* technique, which a dimension of matrix becomes smaller.

## CONCLUSION

In this work, we developed the IPCAPS methodology and motivated its components and assessed its performance via a variety of simulation studies. The simulated datasets used in all experiments were generated using our own tool, called FilestSim. It allows simulating simple and complex data structures with and without outliers. In the majority of the simulations, we compared IPCAPS to other iterative pruning based methods, namely ipPCA, iNJclust, and SHIPS.

## REFERENCES

1. Neuditschko M, Khatkar MS, Raadsma HW. NetView: A High-Definition Network-Visualization Approach to Detect Fine-Scale Population Structures from Genome-Wide Patterns of Variation. Timpson NJ, editor. PLoS ONE. 2012;7:e48375.

2. Price AL, Patterson NJ, Plenge RM, Weinblatt ME, Shadick NA, Reich D. Principal components analysis corrects for stratification in genome-wide association studies. Nat. Genet. 2006;38:904–9.

3. Lawson DJ, Hellenthal G, Myers S, Falush D. Inference of Population Structure using Dense Haplotype Data. Copenhaver GP, editor. PLoS Genet. 2012;8:e1002453.

4. Price AL, Patterson NJ, Plenge RM, Weinblatt ME, Shadick NA, Reich D. Principal components analysis corrects for stratification in genome-wide association studies. Nat. Genet. 2006;38:904–9.

5. Corander J, Marttinen P, Sirén J, Tang J. Enhanced Bayesian modelling in BAPS software for learning genetic structures of populations. BMC Bioinformatics. 2008;9:539.

6. Intarapanich A, Shaw PJ, Assawamakin A, Wangkumhang P, Ngamphiw C, Chaichoompu K, et al. Iterative pruning PCA improves resolution of highly structured populations. BMC Bioinformatics. 2009;10:382.

7. Limpiti T, Intarapanich A, Assawamakin A, Shaw PJ, Wangkumhang P, Piriyapongsa J, et al. Study of large and highly stratified population datasets by combining iterative pruning principal component analysis and structure. BMC Bioinformatics. 2011; 12:255.

8. Chaichoompu K, Abegaz F, Tongsima S, Shaw PJ, Sakuntabhai A, Pereira L, et al. IPCAPS: an R package for iterative pruning to capture population structure. bioRxiv [Internet]. 2017; Available from: http://biorxiv.org/content/early/2017/09/10/186874.abstract

9. Zou F, Lee S, Knowles MR, Wright FA. Quantification of Population Structure Using Correlated SNPs by Shrinkage Principal Components. Hum. Hered. 2010;70:9–22.

10. Clayton D. snpStats: SnpMatrix and XSnpMatrix classes and methods. 2015.

11. Qiu Y, Mei J, details authors of the A library S file A for. rARPACK: Solvers for Large Scale Eigenvalue and SVD Problems [Internet]. 2016. Available from: https://CRAN.R-project.org/package=rARPACK

12. Czarnecki W, Jastrzebski S, Data M, Sieradzki I, Bruno-Kaminski M, Jurek K, et al. gmum.r: GMUM Machine Learning Group Package [Internet]. 2015. Available from: https://CRAN.R-project.org/package=gmum.r

13. Lebret R, Iovleff S, Langrognet F, Biernacki C, Celeux G, Govaert G. Rmixmod: The R Package of the Model-Based Unsupervised, Supervised, and Semi-Supervised Classification Mixmod Library. J. Stat. Softw. [Internet]. 2015 [cited 2016 May 29];67. Available from: http://www.jstatsoft.org/v67/i06/

14. Bodenhofer U, Palme J, Melkonian C, Kothmeier A. apcluster: Affinity Propagation Clustering [Internet]. 2016 [cited 2017 Mar 7]. Available from: https://cran.r-project.org/web/packages/apcluster/index.html

15. Wang MC and D. MeanShift: Clustering via the Mean Shift Algorithm [Internet]. 2016 [cited 2017 Mar 7]. Available from: https://cran.r-project.org/web/packages/MeanShift/index.html

16. Maechler M, Rousseeuw P, Struyf A, Hubert M, Hornik K. cluster: Cluster Analysis Basics and Extensions. 2017.

17. R: Partitioning Around Medoids (PAM) Object [Internet]. [cited 2017 Mar 7]. Available from: https://stat.ethz.ch/R-manual/R-devel/library/cluster/html/pam.object.html

18. Hahsler M, Piekenbrock M. dbscan: Density Based Clustering of Applications with Noise (DBSCAN) and Related Algorithms [Internet]. 2017. Available from: https://CRAN.R-project.org/package=dbscan

19. Sugiyama M, Borgwardt K. Rapid Distance-Based Outlier Detection via Sampling. In: Burges CJC, Bottou L, Welling M, Ghahramani Z, Weinberger KQ, editors. Adv. Neural Inf. Process. Syst. 26 [Internet]. Curran Associates, Inc.; 2013. p. 467–475. Available from: http://papers.nips.cc/paper/5127-rapid-distance-based-outlier-detection-via-sampling.pdf

20. Tian C, Plenge RM, Ransom M, Lee A, Villoslada P, Selmi C, et al. Analysis and Application of European Genetic Substructure Using 300 K SNP Information. PLoS Genet. 2008;4:e4.

21. Heath SC, Gut IG, Brennan P, McKay JD, Bencko V, Fabianova E, et al. Investigation of the fine structure of European populations with applications to disease association studies. Eur. J. Hum. Genet. EJHG. 2008;16:1413–29.

22. Schwarz G. Estimating the Dimension of a Model. Ann. Stat. 1978;6:461–4.

23. Huckins LM, Boraska V, Franklin CS, Floyd JAB, Southam L, GCAN, et al. Using ancestry-informative markers to identify fine structure across 15 populations of European origin. Eur. J. Hum. Genet. EJHG. 2014;22:1190–200.

24. Bhatia G, Patterson N, Sankararaman S, Price AL. Estimating and interpreting FST: The impact of rare variants. Genome Res. 2013;23:1514–21.

25. Browning SR, Browning BL. Haplotype phasing: Existing methods and new developments. Nat. Rev. Genet. 2011;12:703–14.

26. Gibbs RA, Belmont JW, Hardenbol P, Willis TD, Yu F, Yang H, et al. The International HapMap Project. Nature. 2003;426:789–96.

27. A global reference for human genetic variation. - PubMed - NCBI [Internet]. [cited 2017 Aug 2]. Available from: https://www.ncbi.nlm.nih.gov/pubmed/26432245

28. Marchini J, Howie B. Genotype imputation for genome-wide association studies. Nat. Rev. Genet. 2010;11:499–511.

29. Beretta L, Santaniello A. Nearest neighbor imputation algorithms: a critical evaluation. BMC Med. Inform. Decis. Mak. [Internet]. 2016;16. Available from: http://www.ncbi.nlm.nih.gov/pmc/articles/PMC4959387/

30. Purwar A, Singh SK. Hybrid prediction model with missing value imputation for medical data. Expert Syst. Appl. 2015;42:5621–31.

31. Masconi KL, Matsha TE, Erasmus RT, Kengne AP. Effects of Different Missing Data Imputation Techniques on the Performance of Undiagnosed Diabetes Risk Prediction Models in a Mixed-Ancestry Population of South Africa. PLOS ONE. 2015;10:e0139210.

32. Limpiti T, Amornbunchornvej C, Intarapanich A, Assawamakin A, Tongsima S. iNJclust: Iterative Neighbor-Joining Tree Clustering Framework for Inferring Population Structure. IEEE/ACM Trans. Comput. Biol. Bioinform. 2014;11:903–14.

33. Bouaziz M, Paccard C, Guedj M, Ambroise C. SHIPS: Spectral Hierarchical Clustering for the Inference of Population Structure in Genetic Studies. PLOS ONE. 2012;7:e45685.

34. Burgess M, Adar E, Cafarella M. Link-Prediction Enhanced Consensus Clustering for Complex Networks. PLoS ONE [Internet]. 2016;11. Available from: http://www.ncbi.nlm.nih.gov/pmc/articles/PMC4874693/

35. Wilkerson MD, Hayes DN. ConsensusClusterPlus: a class discovery tool with confidence assessments and item tracking. Bioinformatics. 2010;26:1572–3.

36. Li H, Liu Y, Chen W, Jia W, Li B, Xiong J. COCA: Constructing optimal clustering architecture to maximize sensor network lifetime. Comput. Commun. 2013;36:256–68.

37. Willing E-M, Dreyer C, Oosterhout C van. Estimates of Genetic Differentiation Measured by FST Do Not Necessarily Require Large Sample Sizes When Using Many SNP Markers. PLOS ONE. 2012;7:e42649.

38. Steinley D. Properties of the Hubert-Arable Adjusted Rand Index. Psychol. Methods. 2004;9:386–96.

39. Cluster and Outlier Analysis (Anselin Local Moran’s I)—ArcGIS Pro | ArcGIS Desktop [Internet]. [cited 2017 Aug 18]. Available from: http://pro.arcgis.com/en/pro-app/tool-reference/spatial-statistics/cluster-and-outlier-analysis-anselin-local-moran-s.htm

